# Catch me if you can: Species interactions and moon illumination effect on mammals of tropical semi-evergreen forest of Manas National Park, Assam, India

**DOI:** 10.1101/449918

**Authors:** U.M. Bhatt, B. Habib, H.K. Sarma, S.L. Lyngdoh

## Abstract

Species interaction plays a vital role in structuring communities by stimulating behavioral responses in temporal niche affecting the sympatric associations and prey-predator relationships. We studied relative abundance indices (RAI) and activity patterns of each species, temporal overlap between sympatric species, and effects of moon cycle on predator-prey relationships, through camera-trapping in tropical semi-evergreen forests of Manas National Park. A total of 35 species were photo-captured with 16214 independent records over 7337 trap nights. Overall, relatively high number of photographs was obtained for large herbivores (11 species, n=13669), and low number of photographs were recorded for large carnivores (five species, n=657). Activity periods were classified into four categories: diurnal (day-time), nocturnal (night-time), crepuscular (twilight), and cathemeral (day and night time) of which 52% records were found in diurnal period followed by 37% in nocturnal phase whereas only 11% photographs during twilight. Small carnivores were strictly nocturnal (leopard cat and civets) or diurnal (yellow-throated marten and mongooses); whereas large carnivores were cathemeral (tiger, leopard, clouded leopard and Asiatic black bear). Analysis of activity patterns throughout the 24-h cycle revealed a high degree of temporal overlap (>60%) among most of the sympatric species; however, differences in the activity peaks were found between most of the species pairs. Moon phase was classified according to the percentage of visible moon surface as new (0-25%), waxing (25-50%), waning (50-75%) and full moon (75-100%). Moon phase did not have any correlation with activity of large carnivore and large prey. The large carnivore followed the feed and starve pattern of cyclic activity. The activity of small carnivore was influenced negatively by moonlight (_partial correlation_ r = −0.221, p<0.01). The result suggests that large carnivores were active non-differentially across moon phases; however, small carnivores showed significantly high activity in darker nights. These patterns indicate that small predators may differ their activity temporally as an anti-predator strategy or otherwise to increase their foraging efficiency.

## Introduction

Species interactions are one of the most studied areas in community ecology, as interspecific behavior can largely regulate the composition and structure of community assemblages [1]. There are numerous studies about coexistence and resource partitioning between carnivores in tropical forests [2,3], but few focuses on activity patterns and temporal segregation. For carnivores, interspecific interactions are particularly relevant because of their role in the top-down control and also serve as flagship species in the conservation of biodiversity in many terrestrial ecosystems [4]. Though, given the vital role of consumers and through trophic cascades, changes in the environment could promote an increase of medium-sized carnivores or mesopredators, due to top predator removal [5] which can cause substantial changes in the dynamics of interaction among sympatric species [6], with adverse effects on subordinate species. Thus, to minimize risks, subordinate species tend to avoid encounters with dominant species [7], by modifying their activity patterns according to that of the dominant species [8]. Often, the prey tries to avoid the times when predators are more active [9] or segregate in other niche dimensions [10].

Moon cycle is reported to play a significant role in activity changes, and several nocturnal animals can alter their activity in response to moonlight variation [11]. Animals may adapt their schedules throughout the circadian cycle to increase their fitness and allow their mutual co-existence [12]. The dynamics between predators and prey depend on these adaptions too, leading to a balance between their activity patterns [13]. For instance, some mammals, such as rodents [11,14] and bats [15] are known to reduce their activity in brighter nights, allocate it to darker periods of the night [16,17] or seek for covered areas [18,14]. This behavior is thought to be due to an increment of predators hunting success during these nights [15,11,14,19,20]. On the other hand, other species, such as some primates [21] and some nocturnal birds [22] are also known to be more active in brighter nights. This increment of activity may be related to both higher predator awareness and increased food uptake success [21,22,]. Amongst abiotic factors, moon cycle is reported to play an essential role in niche adaptions [11,10,23]. Several nocturnal animals change their activity patterns [17,24] and habitat use [18,14,25] due to moonlight and the level of that response classifies species as lunarphobic [15] or lunarphilic [21]. Many species’ interaction with the lunar cycles remain still unknown, however, and a better perception of the responses of other small and large sized mammals to moonlight is therefore required for a full understanding of its effects.

The tropical forest contains some of the highest levels of species diversity and abundance, but many tropical species are cryptic, shy, and secretive, which makes them notoriously difficult to study and their interactions with one another, remain poorly understood [26]. Recently however with camera-traps, monitoring terrestrial rare, cryptic and secretive species in tropical forests has become effective [27,28]. The technique has improved our ability to study terrestrial movements of Asian tropical forest fauna [29], species diversity [30], the associations among species [31], and their habitats [32]. In addition to recording the presence and abundance data of such taxa, date and time of the captures can help in understanding the activity patterns of carnivores and other mammals [29,33,34].

In the current study, we examine activity rhythms and effect of the moon cycle on mammals in the semi-evergreen forest of Manas National Park, India, using camera traps. Objectives of the study were to: 1) determine relative abundance indices (RAI) and species assemblage of MNP; 2) determine activity periods of each species; 3) quantify temporal overlap patterns between species; and 4) investigate moonlight effect on the activity of sympatric species. Such data can be used to study processes shaping ecological communities, especially whether potentially competing species overlap or avoid each other temporally, and how larger species might influence activity of their smaller cohorts in the same habitat. This information contributes to an understanding of species interactions in tropical forests and should assist in developing more suitable management and conservation strategies for forest communities in the Himalayan foothills.

## Materials and Methods

### Study Area

The study was carried out within the 500 km^2^ of Manas National Park (MNP) (26°35’ - 26°50’ N, 90°45’ - 91°15’ E), a UNESCO World Heritage Site, in the state of Assam, India. Manas lies on the borders of the Indo-Gangetic and Indo-Malayan biogeographical realms on a gentle alluvial slope in the foothills of the Himalayas, where wooded hills give way to grasslands and tropical forest. The elevation ranges between 40-170 m moll with an average of 85 m [35]; the monsoon brings extremely heavy rainfall to this region, reaching up to 3,300 mm annually and temperature ranges between 6-37 ^0^C [35]. The park is home to a variety of important mammal species, including the tiger, pygmy hog, hispid hare and Asian elephant [36] and also it supports 22 of India’s most threatened mammal species, as listed in Schedule-I of the Wildlife (Protection) Act of India [37]. Together with the Royal Manas National Park in Bhutan, the park forms one of the largest areas for conservation significance in South Asia, representing the full range of habitats from the subtropical plains to the alpine zone [38]. MNP acquires a special place from conservation aspect owing to its tropical forests, endemism and a long history of social and political conflict [39]. The national park experienced a fifteen-year-long ethnic and political battle starting in the mid-1980s until fledgling peace was restored in 2003 [40]. The violence during the conflict that followed caused large-scale damages to Manas and left the park vulnerable to logging, local hunting, and poaching of important fauna, causing habitat degradation and rapid loss of wildlife [41,42].

## Methods

### 1. Field sampling design

Data on RAI and species assemblage was collected by deploying camera-traps (n=241) during two sample periods: April 2017 to June 2017 (n=112) and December 2017 to May 2018 (n=129), with the whole area divided into a grid system of size 1×1 sq. km (Fig 1). The cameratrap locations were selected based on the presence of carnivore sign, accessibility, terrain features, animal trails and nallahs (seasonal drainages). At each location, a single Cuddeback-color™ digital camera was set by affixing it to trees at the height of approximately 30-45 cm to above the ground. The cameras were triggered by motion sensor within a range of a conical infrared beam and time lag of approximately 1s between the animal detection. Relative abundance index (RAI) was calculated as the sum of all detections for each species for all camera traps over all days, divided by the total number of camera trap nights, and then multiplied by 100 [43]. To maintain statistical independence and to reduce bias caused by repeated detections of the same species, one record of each species per half an hour per hours per camera-trap site was considered as an independent detection and subsequent records were removed [44]. No bait was used, to avoid disproportionate increases in the frequency of some species [45].

**Fig 1.**
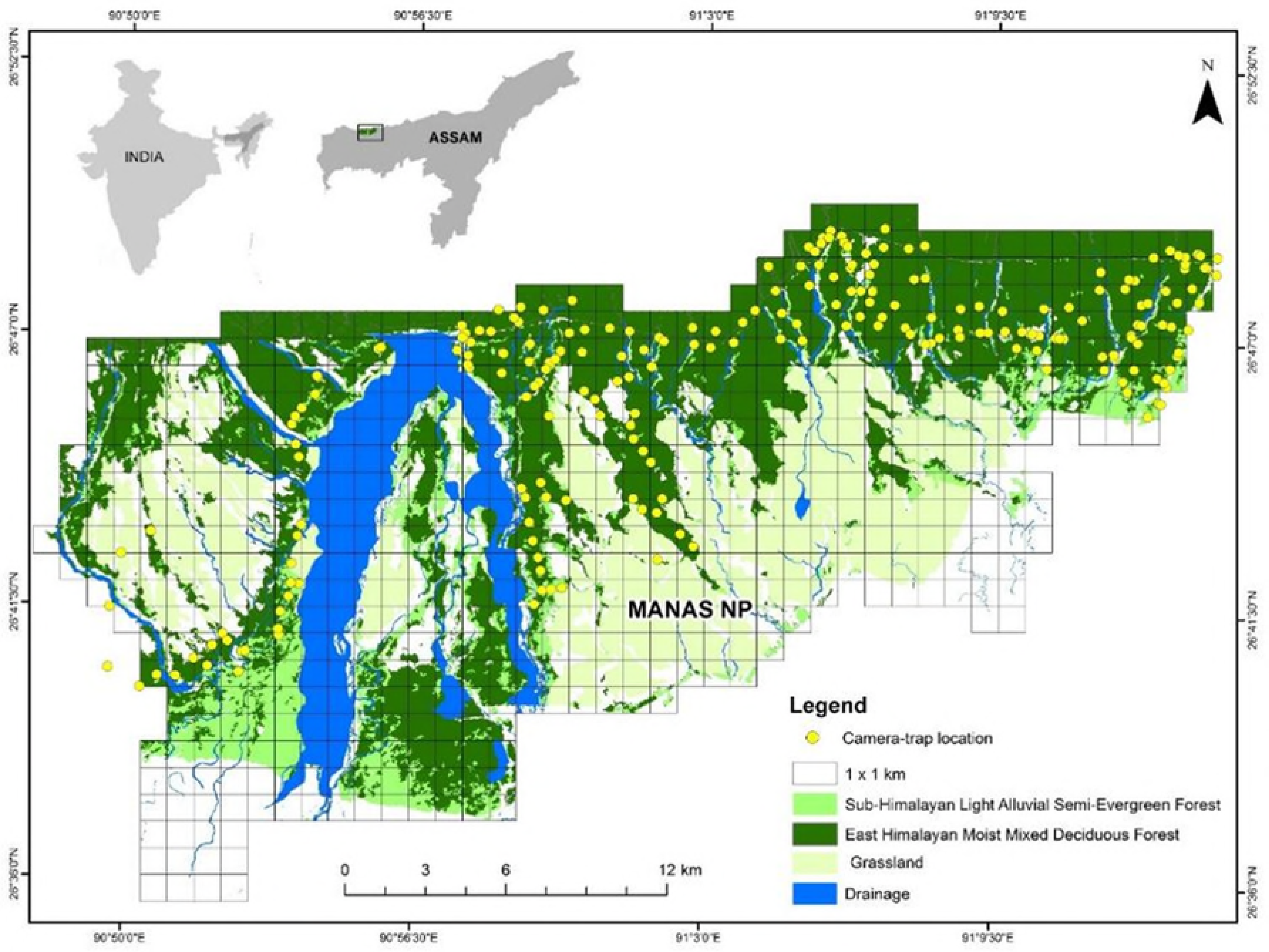
Map of the study area (MNP) showing locations of camera traps (n = 241), grids, drainage and forest cover.

### 2. Activity periods

The date and time printed on the photographs were used to describe diel activity periods of each species. The assumption was made that the number of camera trap records taken at various times is correlated with the daily activity patterns of mammals. The date and time printed on the photographs were used to describe the daily activity patterns of the species. As some species may be partly arboreal, and the camera-traps only recorded activity at ground-level, it is not possible to assess arboreal activity. The observations were classified as diurnal, nocturnal, cathemeral or crepuscular. Photos that captured an hour before and after sunrise and sunset were defined as crepuscular [46]. Sunset and sunrise hours were determined using geographical coordinates of the study area and the Moonphase SH software (version 3.3; Henrik Tingstrom, Kalmar, Sweden). Species were classified as diurnal (<10% of observations in the dark), nocturnal (<90% of observations in the dark), mostly diurnal (between 10-30% of observations in the dark), mostly nocturnal (between 70-90% of observations in the dark) and crepuscular (50% of observations during the crepuscular phase), the rest of the species were classified as cathemeral [47].

### 3. Activity analysis

Kernel density estimation curves were used to describe the activity patterns of each species; a non-parametric way to estimate the probability density function of a distribution of records which assumes that an animal is equally likely to be captured at any time as long as it is active [47]. Overlap coefficients among the daily activity patterns of sympatric carnivores and potential prey were estimated using Overlap package [47] for R-software version 3.1.2 (R Development Core Team, 2011). Overlap coefficients (Δ) is defined as the area under the curve that is formed by taking the minimum of the two density functions at each time point ranging from 0 (no overlap), if species have no common active period, to 1 (complete overlap), if the activity densities of two species are identical [48]. The chosen estimator for overlapping was Δ_1_ or Δ_4_, depending upon the sample size. Δ_4_ estimator for the coefficient of overlap was used if both samples are larger than 50, whereas Δ_1_ was used for small sample size [47]. Data were bootstrapped (99 samples) to extract 95% confidence intervals (CI) from the overlap coefficients [26,49].

### 4. Moon phase

Moonphase SH software, version 3.3 was used to assess the effect of the moon phase on the activity of mammals. The software classifies moon phase of records, according to the percentage of visible moon surface, as follows: 0-25% [New Moon (New)], 25-75% [first quarter - Waxing Moon (Wx) & last quarter - Waning Moon (Wn)] and 75-100% [Full Moon (Full)]. Then, the records from each moon phase were selected to assess the effect of the moonlight and positioning on the time schedules of large – small carnivores and their potential prey, during lunar cycle. One-way analysis of variance (ANOVA), Tukey HSD (honestly significance difference) for Post-Hoc, and partial correlation tests were conducted to measure the degree of association and pairwise comparisons among records of predator-prey in each moon phase.

## Results

### 1. Relative abundance indices & species assemblage

A total of 35 species were recorded with 16,214 independent records over the whole sampling period of 7337 trap nights. The independent records (n) and relative abundance index (RAI) for the photo-captured species varied from species-wise ranging from *Neofelis nebulosa* (n=7, RAI=0.0011) to *Panthera pardus* (n=298, RAI=0.0417) for large – medium carnivores, from *Melogale moschata* (n=1, RAI=0.0001) to *Prionailurus bengalensis* (n=221, RAI=0.0366) for small carnivore, from *Axis axis* (n=1, RAI=0.0003) to *Elephas maximus* (n=4675, RAI=0.5696) for large herbivores, and from *Caprolagus hispidus* (n=1, RAI=0.0002) to *Gallus gallus* (n=574, RAI=0.0853) for small herbivores. The summarised photo captures with RAI of all the species are given in table 1.

**Table 1.**
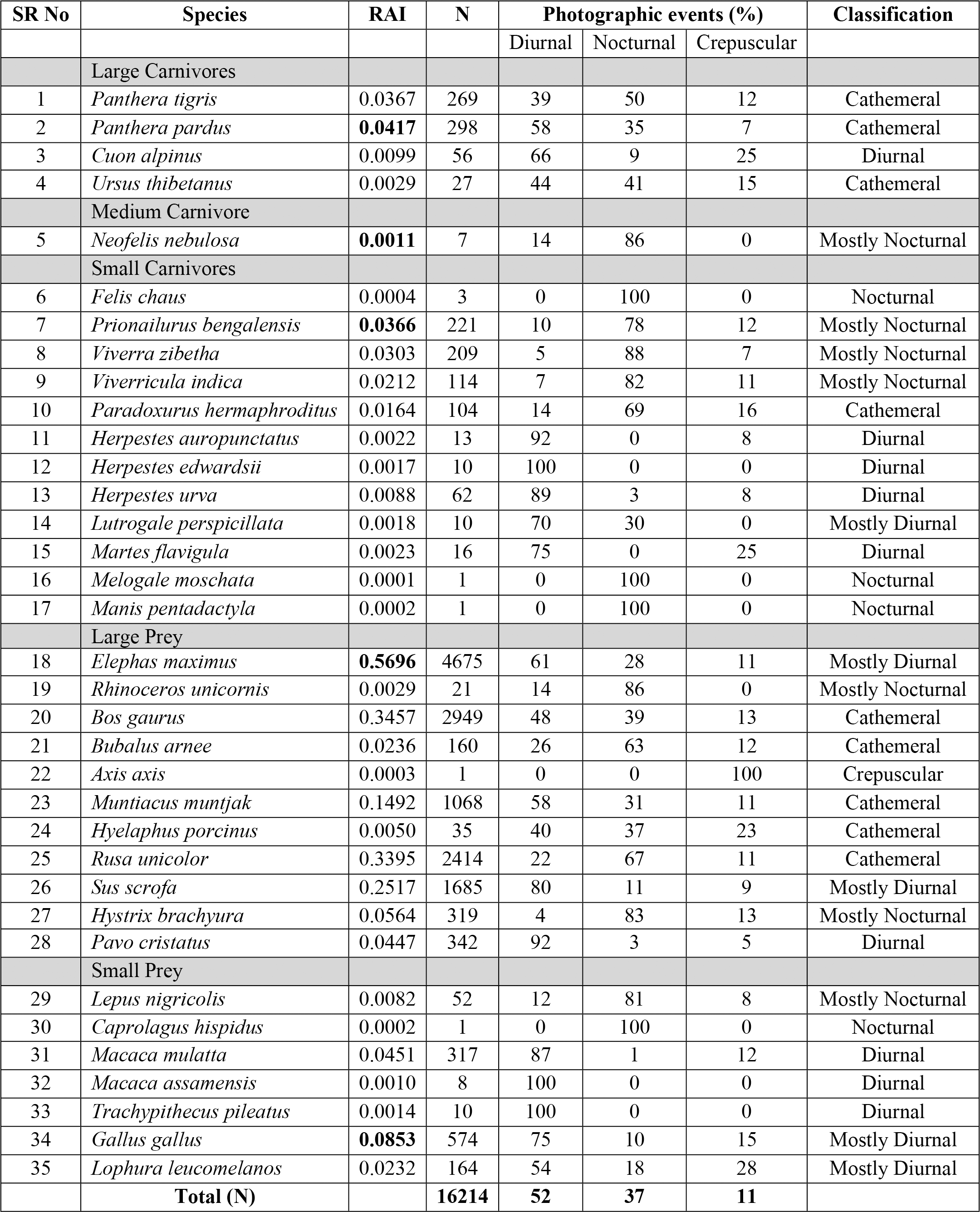
Activity periods of photo-captured species through camera-trapping in Manas

### 2. Activity periods of photo-captured species

Activity periods for 35 species depicts that the boundaries between the categories (diurnal, nocturnal, cathemeral or crepuscular) of activity periods are not sharp (Table 1). However, some animals restrict their routine activities to either the light or the dark phase, and rare observations in the other period may represent activity elicited by unusual circumstances.

#### (i). Carnivores

Small carnivores were either mainly nocturnal such as *Prionailurus bengalensis, Viverra zibetha*, and *Viverricula indica*, or diurnal such as *Martes flavigula, Lutrogale perspicillata, Herpestes urva, Herpestes auropunctatus* and *Herpestes edwardsii*; whereas *Paradoxurus hermaphroditus* with 69% photographs in the dark phase fell under the cathemeral category (Table 1). Out of the five large – medium carnivore species; three large body-sized mammals (*Panthera tigris, Panthera pardus*, and *Ursus thibetanus*) were cathemeral, *Cuon alpinus* was diurnal, and *Neofelis nebulosa* was found with nocturnal nature (Table 1).

#### (ii). Herbivores

Large herbivores were either mainly cathemeral such as *Bos gaurus, Bubalus arnee, Muntiacus muntjak, Hyelaphus porcinus*, and *Rusa unicolor*, or diurnal such as *Elephas maximus*, and *Sus scrofa* (Table 1). The only large herbivore tending toward nocturnality was the *Rhinoceros unicornis* (Table 1). Terrestrial birds (*Gallus gallus, Lophura leucomelanos, Pavo cristatus*) and primates (*Macaca mulatta, Macaca assamensis, Trachypithecus pileatus*) were diurnal; whereas the routine activity of hares (*Lepus nirgicolis, Caprolagus hispidus*) and Himalayan crestless porcupine (*Hystrix brachyura*) suggested nocturnal nature of the species (Table 1).

### 3. Activity pattern and temporal overlap between sympatric species

#### (i). Small carnivores

Eight small carnivores were sufficiently common to evaluate their diel activity pattern (Fig 2). Civets and leopard cats were active during night hours; whereas mongooses and yellow-throated martens were active during the daytime (Fig 2). All four nocturnal small carnivores had shown high temporal overlap between them, with the highest overlap was found between leopard cat and small Indian civet with an overlap coefficient, Δ_4_=0.93 (±0.15), followed by leopard cat and large Indian civet (Δ_4_=0.90); whereas least overlap (Δ_4_=0.74) was found between large Indian civet and palm civet (Fig 2). Leopard cat had a strong bimodal pattern, with a stronger peak at around 23:00 hr and a less pronounced peak from about 01:00 to 04:00 hr. Large Indian civet also showed the bimodal pattern as it increases post-sunset and reaches its peak at around 22:00 hr and then starts to decline; again, it starts rising post-midnight and reaches its peak at about 01:00 hr and then begins to fall (Fig 2). Other two civets (small Indian civet and Asian palm civet) had also shown a bimodal pattern, but with differences in peaks; less pronounced peaks for both the species were between 19:00 to 23:00 hr and 03:00 hr, whereas stronger peaks were about at 04:00 hr and 17:00 to 22:00 hr respectively (Fig 2). High temporal overlap was found between all the four diurnal small carnivores, with the highest coefficient value of Δ_1_=0.84 (±0.09) between crab-eating mongoose and yellow-throated marten, followed by crab-eating mongoose and grey mongoose (Δ_1_=0.77); whereas least coefficient value (Δ_1_=0.54) was found between small Indian mongoose and yellow-throated marten (Fig 2). Small Indian mongoose showed a unimodal pattern, and it increases post 05:00 hr and reaches its peak at around 11:00 hr and then starts to decline gradually until 18:00 hr (Fig 2). Grey mongoose, crab-eating mongoose, and yellow-throated marten were active throughout the light phase, had a bimodal activity pattern, with a stronger peak at 06:00, 15:00 and 16:00 hr, whereas less pronounce peak at 15:00, 08:00 and 06:00 hr respectively (Fig 2). *Melogale moschata* (n=1), *Felis chaus* (n=3), and *Lutrogale perspicillata* (n=10) had the fewest detections and therefore, were not considered for activity analysis (Table 1).

**Fig 2.**
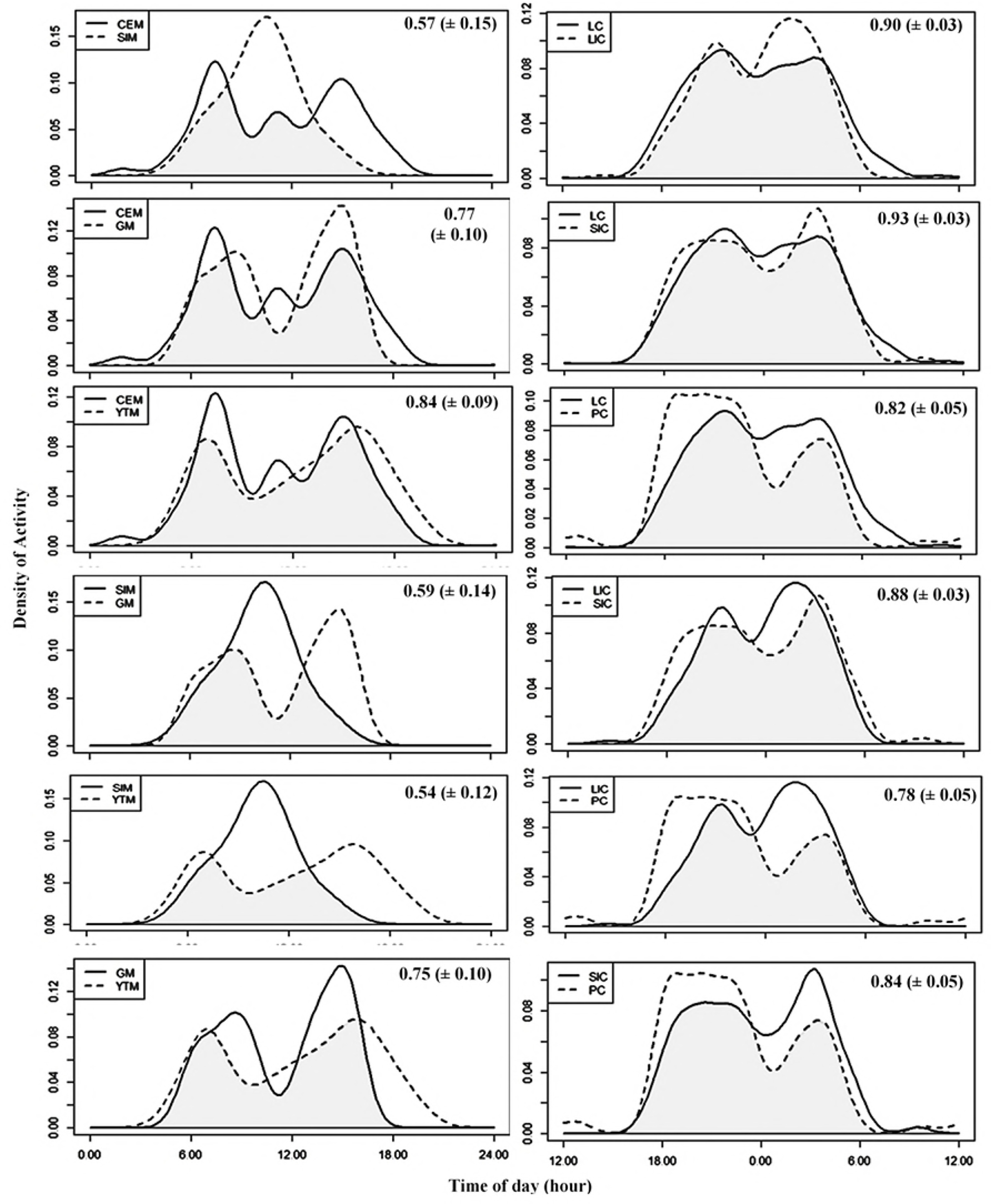
Temporal overlap among small carnivores in Manas National Park, Assam, India. Individual photograph times are indicated by the short vertical lines above the x-axis. The overlap coefficient (Δ_1_ / Δ_4_) is the area under the minimum of the two density estimates, as indicated by the shaded area in each plot. The abbreviations of species’ names are CEM-Crab-eating Mongoose, SIM-Small Indian Mongoose, GM-Grey Mongoose, YTM-Yellow Throated Marten, LC-Leopard Cat, LIC-Large Indian Civet, SIC-Small Indian Civet, and PC- Palm Civet.

#### (ii). Large carnivores

Among the activity patterns of large carnivores, tigers and leopards showed the highest daily activity overlap Δ_4_ = 0.82 (± 0.03) for any 2 species of top carnivores in the study area, followed by leopards and Asiatic black bears (Δ _1_ = 0.82); whereas lowest overlap Δ_1_ = 0.10 (± 0.07) was found between clouded leopards and dholes (Fig 3). Leopard was active throughout the day and night but was more active during daylight, with peaks in the early morning and late afternoon; tiger had also shown cathemeral activity pattern but was least active from about 10:00 to 15:00 hr (Fig 3). Two activity peaks (between 21:00 and 23:00 hr and between 2:00 and 4:00 hr) were observed for clouded leopards, suggesting a bimodal activity pattern of the species (Fig 3). Dholes, showed a unimodal pattern of activity, with peaks between 06:00 to 09:00 hr (Fig 3). However, Asiatic black bears were relatively cathemeral, with 44% of their photo-captures obtained during daytime and 41% records obtained during night-time; whereas least activity was observed between 20:00 to 02:00 hr (Table 1, Fig 3). A synchronized least-active pattern was noted between midnight and 03:00 hr for all the detected large carnivores (Fig 3).

**Fig 3.**
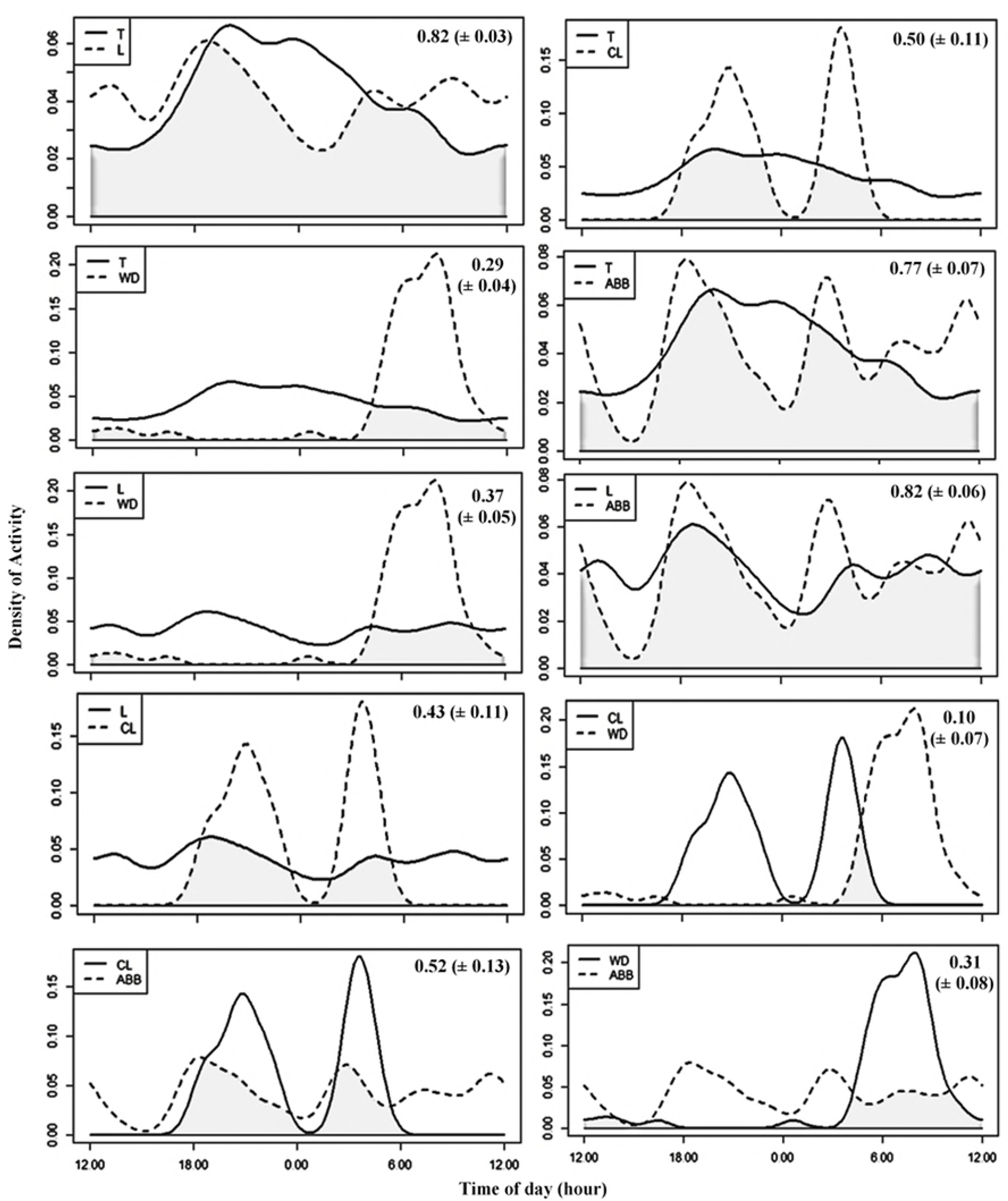
Temporal overlap among large carnivores in Manas National Park, Assam, India. Individual photograph times are indicated by the short vertical lines above the x-axis. The overlap coefficient (Δ_1_ / Δ_4_) is the area under the minimum of the two density estimates, as indicated by the shaded area in each plot. The abbreviations of species’ names are T-Tiger, L-Leopard, CL-Clouded Leopard, WD-Wild Dog, and ABB-Asiatic Black Bear.

#### (iii). Carnivores and their prey

High temporal overlap was found among nocturnal prey species such as Indian hare with leopard cats and civets whereas red junglefowl, and kalij pheasant was active during the daytime, hence had shown large overlaps with mongooses and yellow-throated marten (Fig 4a). Activity patterns of large carnivores and its prey showed variable temporal overlap with the highest overlap between tiger and wild buffalo (84%) followed by sambar (84%), hog deer (84%), and gaur (79%) (Fig 4b). In case of leopard highest overlap was found with hog deer (76%) followed by gaur (74%), barking deer (72%) and wild buffalo (72%) (Fig 4b). Clouded leopard had shown highest overlap with Himalayan crestless porcupine (66%), followed by sambar (65%) whereas dhole had maximum overlap with wild boar (52%) and barking deer (50%) (Fig 4b). Chital (n=1) and hispid hare (n=1) had only one detection and therefore, were not considered for the analysis (Table 1).

**Fig 4.**
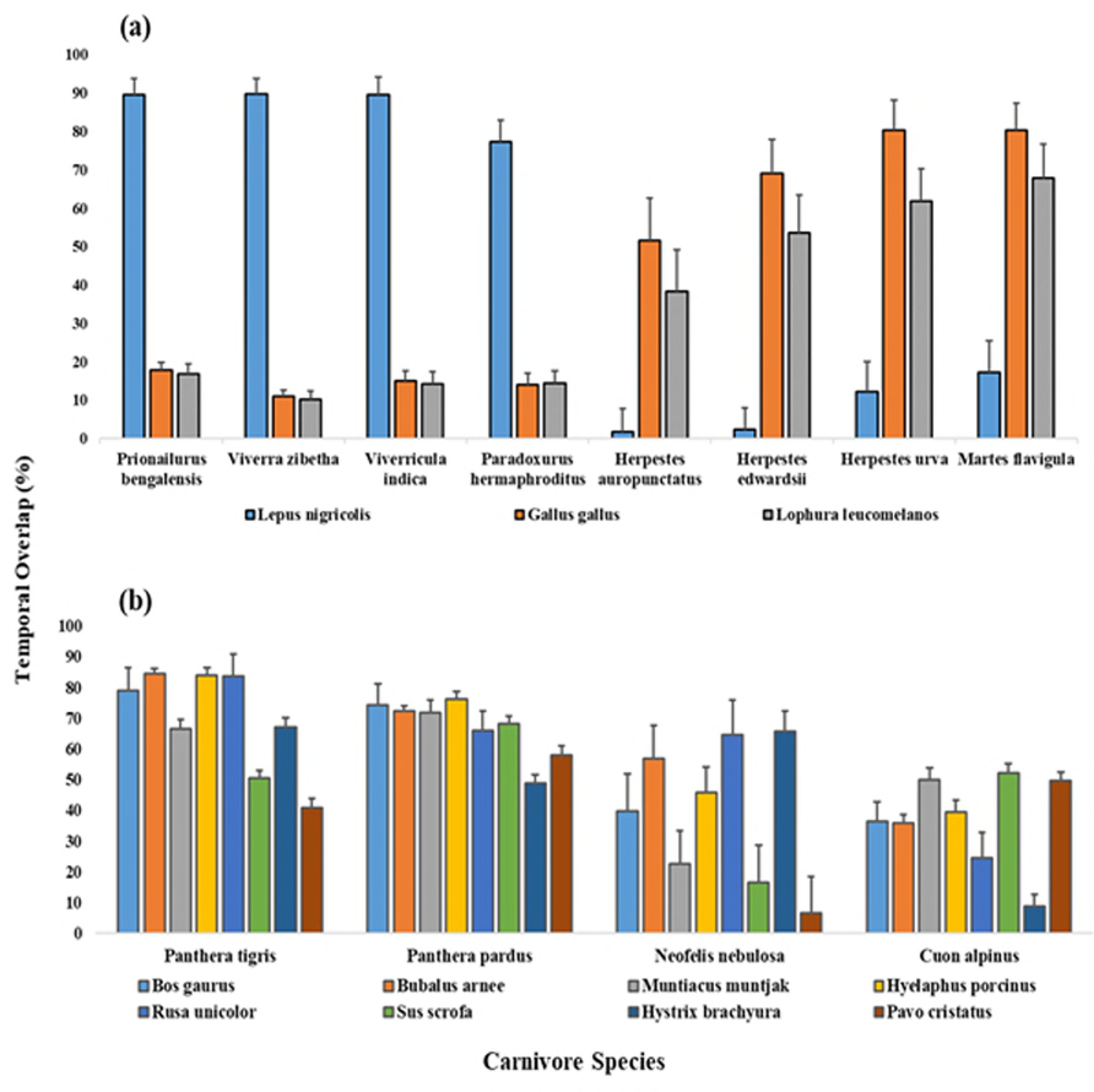
Pairwise temporal overlap (Δ_1_ / Δ_4_) between (a) small carnivore vs. prey and (b) large carnivore vs. prey in Manas National Park, Assam, India. The bars indicate percentage temporal overlap among carnivore species with potential prey and the whiskers above the bars indicate standard errors.

### 4. Moon phase effect on prey-predator relationship

Photographs of large carnivores, small carnivores, and potential prey species were analysed during four moon cycles (New, Wx, Full, and Wn) (Fig 5). Differences were observed in carnivore community with respect to the moon cycle, with highest records of large carnivores in full moons except for dhole while small carnivores had more photographs at new moon phase, except for Asian palm civet (Fig 5). Dhole activity was found mainly diurnal with only 9% photographs in nocturnal periods, out of which around 58% records were recorded in darker nights (Fig 5). On the other hand, Asian palm civet had more photographs in full moon (33%); and 21% photographs were recorded in new moon phase (Fig 5). All three photo-captured small prey such as red junglefowl, kalij pheasant, and Indian hare had more photographs in the new moon phase (Fig 5). Larger prey showed almost uniform activity in all moon phases, with highest records in new moon (wild buffalo, hog deer, wild boar, Himalayan crestless porcupine and Indian peafowl); whereas the remaining three species (gaur, barking deer and sambar) had more photographs in a full moon (Fig 5).

**Fig 5.**
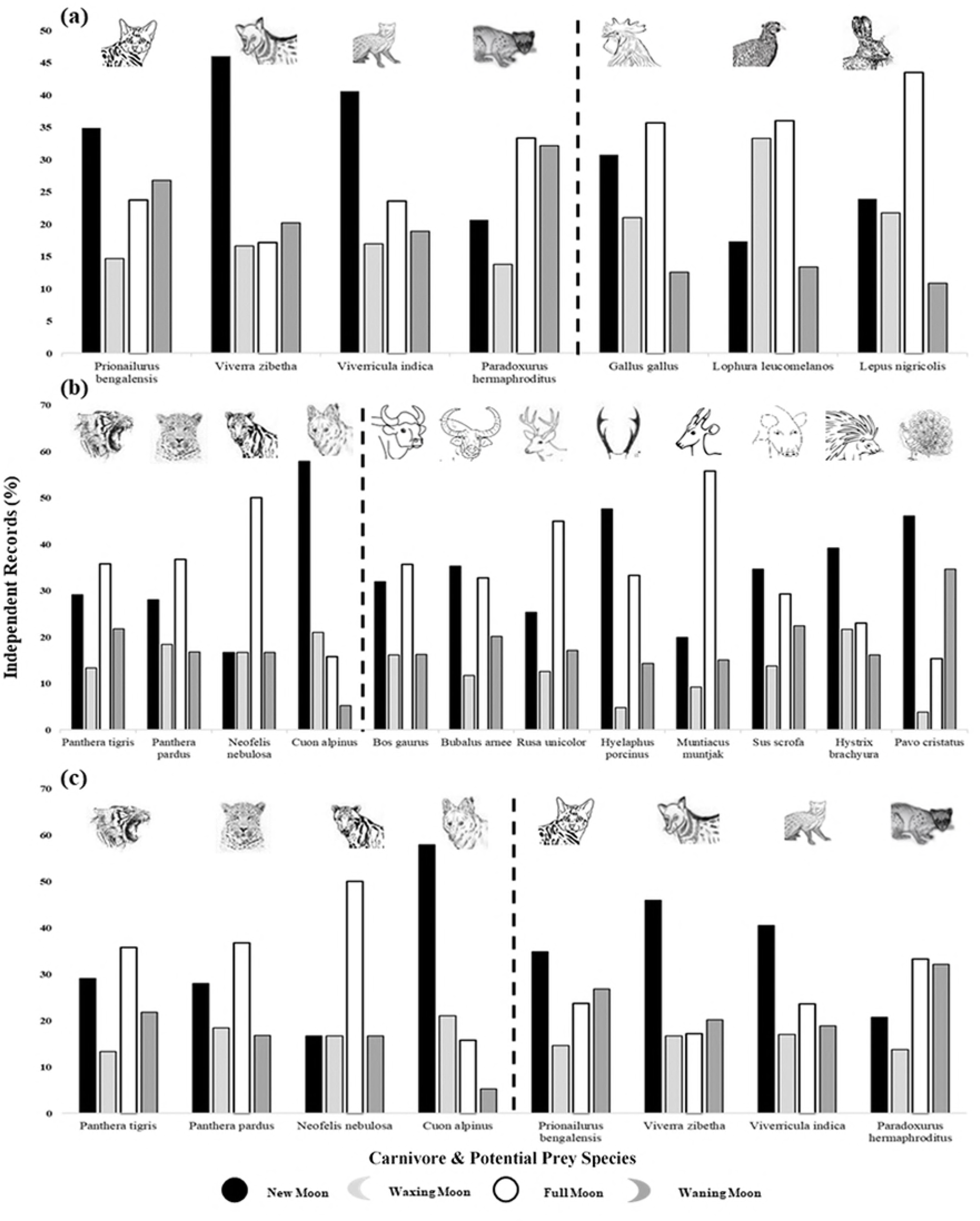
The proportion of nocturnal records of (a) small carnivore vs. prey, (b) large carnivore vs. prey, and (c) large carnivore vs. small carnivore in different moon phases in Manas National Park, Assam, India. The bars indicate species records in different moon phases. The dashed line is to separate small carnivore and their prey, large carnivore and their prey, large carnivore and small carnivore respectively.

One-way ANOVA result pointed significant difference only for small carnivore (F= 5.007, p<0.005) and for small prey (F= 3.697, p<0.05). In case of small carnivores, the Tukey’s HSD for post-hoc result showed significant more records in new moon (mean differences= 1.52, p<0.005) and waning moon (mean differences= 1.38, p<0.05) than in full moon. For small prey, more photo-captures were recorded in waxing moon (mean differences= 1.21, 1.49; p<0.05) than in new moon and waning moon respectively.

The results of the partial correlation depicted a negative relation between small carnivore and moon visible surface while controlling for small prey (r= −0.221, p<0.001) and large carnivore (r= −0.213, p<0.01) (Table 2). However, Pearson’s product-moment correlation also known as the zero-order correlation showed statistically significant, negative correlation between small carnivore and moon visible surface (r= −0.205, p<0.01), without controlling for small prey or large carnivore. This suggests that small prey or large carnivore had very little influence in controlling the relationship between small carnivore and moon visible surface.

**Table 2.**
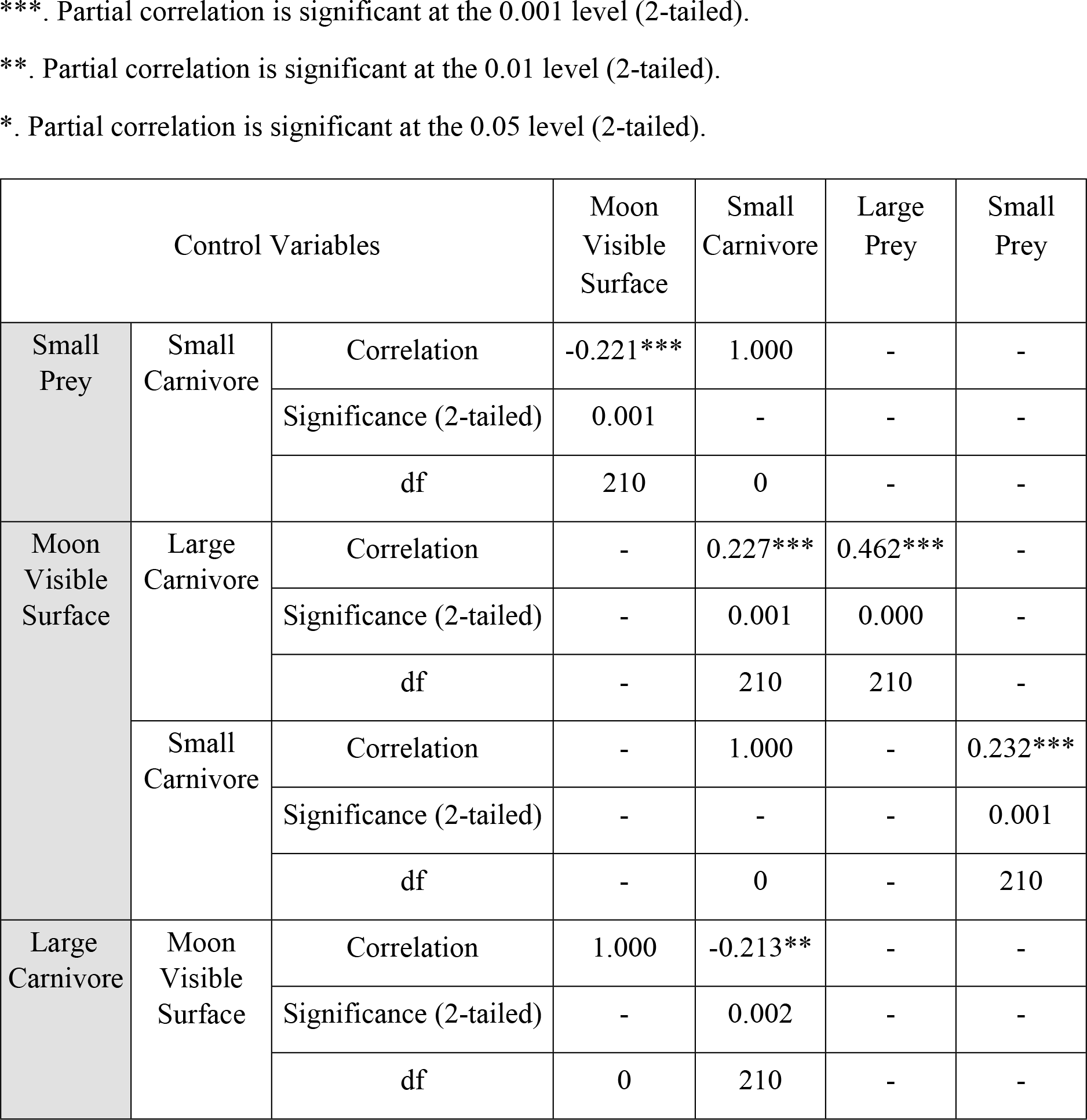
Partial correlation test (r) for degree of association between prey-predator and moon visible surface using different control variables in Manas National Park, Assam, India.

## Discussion

The current study provides baseline information on activity patterns and temporal overlaps of mammals of Manas National Park as well as it is also the first of its kind of research on moon illumination and effect of moon phases on prey-predator interactions in tropical forests of India. Results from the present study are mainly concordant with basic accounts of natural history (i.e., whether a species is most active during the day or night) [50]. We also compared our results with those of previous studies on species body size, activity pattern and temporal overlaps of mammalian fauna. The lunar cycle results largely showed that the moonlight has a stronger effect on the activity of the prey than on the behavior of the predator.

Evaluation of the camera trapping data revealed that the study area had a healthy habitat for the mammalian fauna. All the major fauna from MNP was photo-captured during the survey confirming 35 species. Out of the 35 recorded species, 1 (Chinese pangolin) is classified as Critically Endangered, 6 (tiger, dhole, Asiatic elephant, wild water buffalo, hog deer and hispid hare) are classified as Endangered, 8 (leopard, clouded leopard, Himalayan black bear, one-horned rhinoceros, gaur, sambar and capped langur) are classified as Vulnerable, 1 (Assamese macaque) is classified as Near Threatened while the remaining 19 species are classified as least concern [51]. The previous camera-trapping studies in Manas National Park provides information on relative abundances of tigers and their prey [52], carnivore diversity [53], and density estimation of carnivores and herbivores [40]. Recently, Borah et al. [54] provide info on density estimation of common leopard and clouded leopard; whereas Lahkar et al. [55] explained about diversity, distribution and photo-capture rate of mammals of MNP. In the present study, we examine activity rhythms and the lunar cycle effect on the mammalian fauna in the semi-evergreen forest of Manas National Park.

### 1. Diel activity patterns and temporal overlap

Van Schaik and Griffiths [29] explained variation in activity periods for Indonesian rainforest mammals using species body size as the primary factor influencing activity patterns. The theory suggests that smaller mammals (<10 kg) tend to be specifically nocturnal or diurnal as an anti-predation strategy, whereas larger mammals (>10 kg) are more cathemeral because of energy requirements and associated feeding commitments. The intensive camera-trap survey provided one of the most detailed studies of activity periods in mammals of MNP under natural conditions and classified activity patterns into four categories [29]. In the present study, all the photo-captured small mammals are found to be mainly nocturnal (jungle cats, leopard cats, and civets) or diurnal (smooth-coated otter, yellow-throated marten, and mongooses), as predicted by the van Schaik and Griffiths model. The results showed that the medium-sized mammals are cathemeral (barking deer, hog deer, and wild boar) and diurnal (wild dog), and the larger-sized mammals such as tiger, leopard, Asiatic black bear, gaur, wild buffalo, and sambar are active during both day and night hours which is also in accordance with the Schaik and Griffiths model.

We found differences in the activity peaks of tiger and leopard, but there was no active temporal separation between predators probably owing to their similar morphology and hunting strategies [2]; however, significant time overlap between them was evident. Tigers are opportunistic predators [56] and had considerably higher activity overlap (>75%) with gaur, wild buffalo, sambar and hog deer [57,58]. However, their diet includes birds, fish, rodents, insects, amphibians, reptiles in addition to other mammals such as primates and porcupines [56]. The present study also found higher overlap (>65%) with Himalayan crestless porcupine as compared to leopard’s overlap. Leopard’s activity overlapped (>70%) with all the prey species ranging from medium to large sized prey [59,60]. Leopard showed higher temporal overlaps with medium-sized prey [60] such as barking deer in comparison with tiger’s overlap. Asiatic black bear tends to be diurnally active [61] or crepuscular [62]. The current study found cathemeral nature of the species as it has to spend most of its time in search of food for energy requirements under a very high competition with conspecifics (for vegetal food/and animal matter) as well as other carnivores (for the animal matter) [63].

The study showed that clouded leopard activity was predominantly nocturnal, similar to the studies of Gumal et al. [64], and Azlan & Sharma [65] from Peninsular Malaysia, and also Kanchanasaka [66], and Grassman et al. [67] from Southern Thailand. Austin et al. [68] recorded activity peaks at crepuscular hours in two radio-collared clouded leopards. However, the overall activity pattern from radio-telemetry studies (n=4) indicated two peaks at 18:00-02:00 hr and 08:00-12:00 hr [69]. The present study also found a bimodal pattern but with different peaks at 21:00-23:00 hr and 2:00-4:00 hr. In case of its prey, the study found the high temporal overlap with sambar and Himalayan crestless porcupine [57] as compared to the other prey species. It is possible that clouded leopard terrestrial activity is higher at night-time due to the avoidance of leopards in the study area being more active on trees during daytime (A. Wilting, pers. comm). However, studies suggest clouded leopards be more terrestrial [70,71,72] with the use of trees primarily for resting [70,73]. The low capture rate of 7 photos in 7337 trap nights in our study, does not necessarily reflect low numbers of the felid, but rather a decreased probability to capture it along wildlife trails and roads that are frequented by high numbers of leopards and tigers, the top predator of the area. The species is known to use a dimension that was not covered in our sampling, namely trees higher up than 60 cm above ground [74].

The only canid species recorded during the study was wild dog (dhole). Dholes showed less temporal overlap with their dominant competitors (tiger, leopard and clouded leopard) because they were more active during the daytime and crepuscular hours and less active in full darkness, similar to most other studies of India and Southeast Asia [67,75,76,77]. Dholes’ diet includes a wide variety of prey species, ranging from small rodents and hares to gaur [2,78,76,79]. In tropical semi-evergreen forests of Southeast Asia, the species appear to persist in smaller packs and consume medium-sized prey [75], as smaller packs are more energetically advantageous in the rainforest where large prey species are scarce, thick vegetation favors stalk and ambush hunting techniques over cursorial hunting, and competition with tiger and leopards [80]. The present study also found the high temporal overlap of dhole with medium-sized prey such as barking deer and wild boar as compared to other large-sized prey.

The present study recorded the two small cats (leopard cat and jungle cat) to be strictly nocturnal which is consistent with other reported studies only in case of leopard cat [81,82], yet there are studies contradicting nocturnality of leopard cats [83,65,84]. According to Prater [81], jungle cat is diurnal as well as crepuscular and can even kill porcupine species which are nocturnal. In this study, jungle cat is found to be strictly nocturnal as reported by Majumder et al. [85], though our result is insignificant because of only three captures of a jungle cat. The small Indian civets and large Indian civets are found active in nocturnal hours which is consistent with other reported studies [86,87]; whereas with 69% photographs in darker hours of Asian palm civet also supports other study of Duckworth [88] and Azlan [89] which suggests crepuscular or nocturnal nature of the species. Mongooses are predominantly diurnal or cathemeral [90] and the present study found a strictly diurnal pattern for all the three photo-captured mongooses. The yellow-throated marten was completely a diurnal species in the study area [91] but can hunt both by day and night and also known to attack young deer species [81]. On the other hand, results of the current study showed a Himalayan crestless porcupine as a nocturnal species which is consistent with the previous studies by Menon [92]. Indian hare which is mainly active during crepuscular and nocturnal hours [93] are found to be active only in nocturnal phase in this study. According to several studies [13,94], the diel activity of many felids is associated with the activity pattern of their prey. The main reason of these small cats being nocturnal in the study area could be that rodents (Himalayan crestless porcupine) and hares (Indian hare and hispid hare), their primary preys, are generally nocturnal [81,94] although we could not quantify this point through camera trapping for small-sized prey.

### 2. Do species respond differently to moonlight?

Moonlight has usually been thought to increase predation risk by enhancing the ability of predators to detect prey [14,95], therefore leading to decreased activity or shifts in prey foraging efficiency in the presence of bright moonlight [16,96,97]. The results of the present study demonstrate that the effects of moon illumination on activity across nocturnal mammal species. The response of nocturnal mammals to the moonlight differs among taxa and may vary according to several determinants, such as phylogeny, trophic level, sensory systems and habitat type [13,97]. The current study suggests that the moon phases are also likely to influence how prey distribute their activities through time to face different predation risk periods.

Large-sized prey species activity were not significantly affected by moon phases as they showed uniform activity with highest record of photographs in new moon (wild buffalo, hog deer, wild boar, Himalayan crestless porcupine and Indian peafowl) and full moon (gaur, barking deer and sambar) (Figs 5 and 6b). Large carnivore did not get influenced by moonlight as they follow the feeding and starvation pattern of cyclic activity across a lunar cycle (Fig 6a). We, therefore, suggest that large carnivores switch their type of prey, they hunt in different moon phases, their hunting efficiency increases in the full moon and the greater foraging benefits they take in brighter nights as they are more photo-captured during a full moon (Fig 5).

Small-sized prey species were more active during brighter nights to avoid predation risk against smaller carnivores (Fig 5). This anti-predator behavior is already well recognised for the species such as marsupials and rodents [33]. However, statistics showed that moonlight did not influence the activity of small prey (Fig 6d). Small carnivores displayed a higher level of activity during the darker nights when reduced brightness hampers their visual detections by large carnivores which were active more in brighter nights (Fig 5). Tukey and partial correlation tests also highlighted that moonlight had negative influence on the small carnivore activity; their activity decreases with an increasing moonlight intensity (Fig 6c, Table 2).

**Fig 6.**
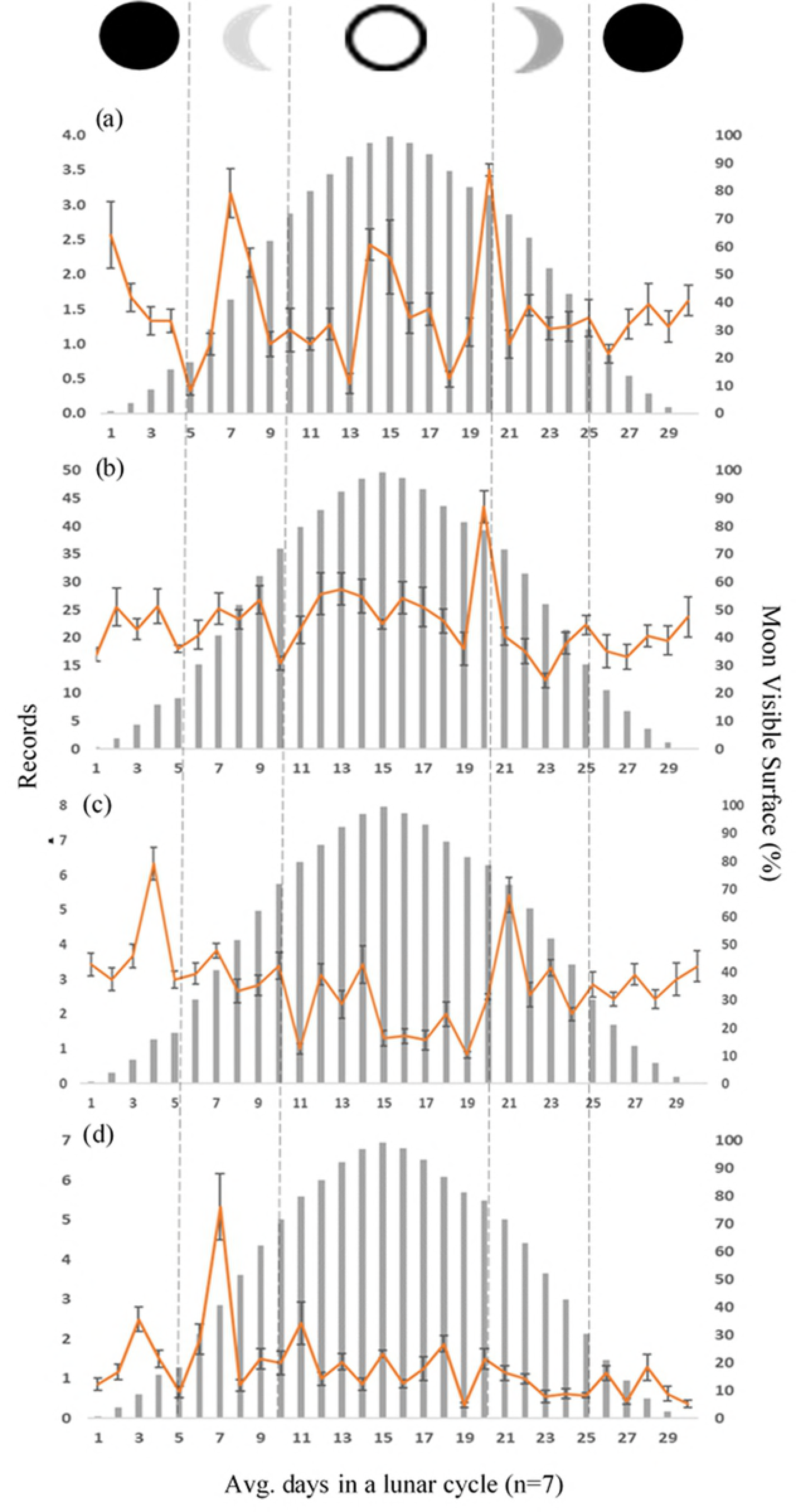
Photo-captures of (a) large carnivore, (b) large prey, (c) small carnivore, and (d) small prey in a lunar cycle in Manas National Park, Assam, India. The bars indicate percentage moon visible surface in a complete lunar cycle. The lines indicate average records in each day from all 7 lunar cycles, and the whiskers above and below the lines indicate standard errors.

## Conclusion & limitation

Our result suggests that despite historical ethnopolitical conflict and continued threats in some areas, MNP supports a diversity of mammalian fauna of conservation concern, including clouded leopards, dholes, tigers and other species. Adaptations are bidirectional and take place over at least two dimensions: spatial and temporal [9,98] and our study focuses primarily on the temporal component and provides some interesting insights into the diel activity patterns and temporal overlap among mammals of MNP. The current study also highlights the significance of incorporating moon illumination into movement and activity pattern of mammals as well as interactions between prey-predator in tropical forests of India. The brighter hours or full moon lights shows an inverse relation in the activity pattern of prey and predator. Camera trapping is effective in recording species interaction but with certain limitations such as the inability to account for detection probability, which is bound to vary with species [76]. Placement of camera traps should be done depending on size, habitat and activity pattern of species. Like in our study, some of the species are at least partially, or even predominantly, arboreal such as clouded leopard, small prey and primate species. Hence, activity patterns of such species would be better explained if camera-traps were deployed in species-specific habitats. Moonlight effects were not only related to the trophic level and were better explained by phylogenetic relatedness, visual acuity, and habitat cover.

## Acknowledgements

We thank the director, dean & research co-ordinator, wildlife institute of India. We are thankful to the Doyil Vengayil & Syed Asrafuzzaman, Department of Science and Technology, Government of India to carry out the study on the clouded leopard (*Neofelis nebulosa*). Paniram, Tapan, Anukul, Dipul, Dipen and Dilli are thanked for their assistance in the field. We thank the Dept., Environment & Forests, Govt., of Assam, field staff of Manas National Park, for permissions and field support.

